# Time spent in conversation over meals predicts default network function: Evidence from a passive mobile-sensing and fMRI study

**DOI:** 10.1101/2024.09.04.611232

**Authors:** Dhaval Bhatt, Jeremy F. Huckins, Subigya Nepal, Andrew T. Campbell, Meghan L. Meyer

## Abstract

Sharing meals with others, often referred to as ‘commensality,’ fosters social connection and may have been essential to human brain evolution. Here, we investigate whether shared meals impact social brain processes through the conversations they afford. If so, then time spent in conversation over meals should affect the default network– a set of interconnected cortical regions reliably associated with the social exchange of ideas. To test this possibility, we combined passive-mobile sensing with neuroimaging. Undergraduate participants had an application on their smartphone for two months that unobtrusively captured their time spent in conversation, as well as their location on and around campus. In between these two months, participants completed a resting state scan. Time spent in conversations around meals (e.g., at cafeterias and restaurants) during the prior month predicted greater functional connectivity in the dorsomedial default network subsystem, specifically between the left inferior frontal gyrus (LIFG) portion and the rest of the subsystem regions. This relationship was preferential to: (1) time spent in conversation over the past (vs. future) month, (2) connectivity with the LIFG (vs. other dorsomedial subsystem nodes), (3) the dorsomedial subsystem (vs. other default network subsystems), (4) eateries (vs. other locations), and (5) time spent *around conversation* at eateries, rather than time at eateries more generally. Follow-up analyses further revealed results were driven by a ventral portion of the LIFG, with peaks in voxel-connectivity associated with language, social, memory, and affective processes, as determined by the meta-analytic platform, NeuroSynth. Time spent in conversation over meals may exercise social cognitive processes supported by the dorsomedial subsystem, a network key to social communication, understanding, and connection.

## Introduction

Eating together–referred to as ‘commensality’ –is a constant across human history, cultures, and lifespans (Dunbar, 2017; Powdermaker, 1932; Rozin, 2014). Commensality is unlikely coincidental, and instead may help explain human brain evolution. One hypothesis of human brain evolution is that the advent of cooking generated more calorically dense meals, providing the additional nutrients needed to expand the human cortex (Herculano-Houzel, 2017; Wrangham, 2009). Another hypothesis is that the social cognitive demands of group living–the need to keep track of friends and enemies in a social network–drove human brain-to-body size ratio (Shultz & Dunbar, 2007). Commensality connects these hypotheses: shared meals provide calories and social interaction. Yet, despite possibly guiding human brain evolution, whether and how commensality in contemporary life impacts brain function remains unknown.

Commensality may impact social brain mechanisms through the conversations they afford. Sharing a meal is an opportunity to exchange ideas and compare notes on everyday life. People rarely eat together in silence and conversations are a primary way humans maintain social bonds (Dunbar, 1996, 2004; Mastroianni et al., 2021). In naturalistic conversations, the dominant information exchanged is about people–either speakers sharing information about themselves or other people (Dahmardeh & Dunbar, 2017; Dunbar, 2017; Dunbar et al., 1997; Emler, 1992; Mesoudi et al., 2006). One overlooked feature of this prior work is that a lot of the data was collected from conversations around meals, for example while talking in cafeterias (Dahmardeh & Dunbar, 2017; Dunbar et al., 1997). Shared meals may be a salient context to share social knowledge through conversation, perhaps impacting social brain mechanisms.

The brain’s default network–a set of interconnected cortical regions–is reliably associated with many social cognitive processes, including communicating our ideas to others and listening to others share their ideas with us (Baek & Falk, 2018; Collier & Meyer, 2020; Zadbood et al., 2017). The default network can be subdivided into three subsystems: the dorsomedial (dMPFC) subsystem, core subsystem, and medial temporal lobe (MTL) subsystem (Andrews-Hanna et al., 2010; Yeo et al., 2011). Of these subsystems, regions in the dMPFC subsystem are the most preferentially associated with high-level social cognition, such as inferring mental states, retrieving social semantic knowledge, and reasoning about interpersonal relationships (Andrews-Hanna et al., 2010; Buckner et al., 1999; Collier & Meyer, 2020; Mitchell et al., 2004; Satpute et al., 2014; Saxe & Kanwisher, 2003; Van Overwalle & Baetens, 2009). For example, the temporoparietal junction (TPJ) region in this network plays a causal role in mental state inference (Bardi et al., 2017; Young et al., 2010) and the dMPFC and superior temporal sulcus (STS) are selectively responsive to real-world social interactions (Lee Masson et al., 2024; Wagner et al., 2016). Also relevant to conversation, another region frequently included in this subsystem (Yeo et al., 2011)– the inferior frontal gyrus–is implicated in both language processing (Klaus & Hartwigsen, 2019; Uddén & Bahlmann, 2012; Xu et al., 2020) and inhibiting our own point-of-view to understand others’ perspectives (Lamm et al., 2010; Morawetz et al., 2022; van der Meer et al., 2011). If shared meals afford the social exchange of ideas through conversation, then more time engaged in them should relate to the dorsomedial default network subsystem.

One difficulty in testing this prediction is that sharing a meal is not particularly conducive to the brain imaging environment. To date, undergoing brain imaging requires participants to be isolated in a large machine while lying on their back. Although food can be delivered to participants during a scanning session (Babbs et al., 2013; Kroemer et al., 2016), the experience of a shared meal is difficult to capture. In fact, even outside of the constraints of MRI scanning, objectively measuring real-world commensality, and the role of conversation in it, is challenging. For example, self-report assessments of time spent in conversation over meals are likely prone to retrospective errors (Collopy, 1996; Hirschi et al., 2023; Mastrandrea et al., 2015; Mastroianni et al., 2021).

To overcome these methodological barriers, we combined brain imaging methods with passive mobile-sensing to unobtrusively measure time spent in real-world conversations around meals. Prior work has successfully linked passive-mobile sensing with brain imaging approaches (Heller et al., 2020; Huckins et al., 2019; Obuchi et al., 2020). For example, data from cell phone global positioning systems (GPS) has been used to show that the positive affect generated by visiting diverse locations predicts resting state functional connectivity between brain regions implicated in reward and spatial navigation (Heller et al., 2020). Prior work has also used cell phone microphones to passively quantify face-to-face conversation and found that conversations predict resting state functional connectivity between the amygdala and ventromedial prefrontal cortex (Obuchi et al., 2020), a circuit implicated in many affective processes (Jalbrzikowski et al., 2017; Kim et al., 2011; Motzkin et al., 2015; Tottenham & Galván, 2016). We combined aspects of each of these approaches – (1) GPS coordinates and (2) time spent in conversation – to test whether time spent in conversations around meals relates to resting state connectivity in the dorsomedial subsystem of the brain’s default network.

To test our ideas, we capitalized on an existing dataset in which first-year college students had a smartphone application on their cell phones– the StudentLife App (Wang et al., 2014, 2018) –that passively tracked their GPS coordinates and time spent in face-to-face conversations for roughly two months. Because first-year students attending this university are required to live in dormitories on campus, they eat most of their meals in college cafeterias and local cafes and restaurants–hereafter referred to as “eateries.” This allowed us to compute, for each participant, the amount of time they spent in conversations in eateries over the course of two months. We also computed time spent in conversations at other locations, such as classrooms and student housing, which allowed us to investigate the extent to which our findings were specific to conversations around meals, above and beyond time spent in conversation more generally.

In between the two months of passive mobile sensing data collection, participants completed a functional magnetic resonance imaging (fMRI) scanning session in which they underwent a standard resting state scan designed to unobtrusively measure brain function. The scanning session occurred in the middle of mobile sensing data collection. This allowed us to arbitrate between two potential ways commensality may relate to dorsomedial subsystem function. One possibility is that there is a trait-like relationship: individuals who spend more time in conversation around meals may also have greater dorsomedial subsystem connectivity. Many stable traits have been linked to default network function (Vaidya & Gordon, 2013). For example, in the social domain, individuals with stronger dorsomedial subsystem resting state functional connectivity score higher on trait measures of prosocial behavior (Inagaki & Meyer, 2020). Similarly, dorsomedial subsystem connectivity may relate to “trait like” commensal behavior, with time spent in conversation around meals across the months of passive mobile sensing data collection linked to the integrity of this subsystem.

Alternatively, time spent in conversations around meals during the past (vs. future) month may preferentially impact dorsomedial subsystem functional connectivity. To the extent shared meals provide opportunities to engage in social cognition, they may exercise the dorsomedial subsystem and enhance its connectivity. Indeed, prior work in humans and non-human primates suggests that changes to the social environment that increase the need for social cognition impact default network function (Gee et al., 2013; Martínez-García et al., 2023; Paternina-Die et al., 2020; Sallet et al., 2011; Tottenham & Galván, 2016). Experimental work also finds increasing social cognition enhances subsequent default network resting state functional connectivity (Collier & Meyer, 2020; Inagaki & Meyer, 2020; Lieberman et al., 2019; Meyer et al., 2015, 2019; Meyer & Collier, 2020; Sippel et al., 2021). The relationship between conversations over meals and default network function may operate most strongly in one direction, with the amount of time spent in conversations around meals during the past (vs. future) month preferentially enhancing dorsomedial subsystem functional connectivity.

## Methods

### Participants

113 first-year college students between the ages of 18-22 years agreed to participate in this study. Participants with irregular fMRI scans (incomplete or short scans; N=5), irregular mobile sensing data (e.g. paused location-tracking; N=8), and missing mobile data (data tracking switched off; N=12) were excluded. This resulted in 88 participants (ages = 18.25 M ± 0.64 SD; 70.5% Females; 60.4% White, 23.4% Asian, 2.6% Black, 10.2% Multi-racial, and 3.4% did not report) with 8 weeks of mobile-sensing data and a resting-state fMRI scan, collected during the academic year. The full study protocol was approved by the Dartmouth College Institutional Review Board (IRB) and participants provided informed consent. Of the 88 participants with usable data, outliers more than 2.5 standard deviations outside of a variable mean were removed from analyses assessing those variables. This included subjects with functional connectivity (N=2) and conversation variables (e.g., total duration spent around conversations in the prior (N=11) and future (N=15) month). We therefore report varying degrees of freedom across results due to missing or drops in conversation data. We picked a 2.5 SD threshold to be somewhat conservative in outlier determination. That said, it is important to note that our results do not depend on this threshold. When a less conservative outlier threshold (i.e., SD > 3.5) is used, the primary results reported below persist: conversation duration over the month before the scan day is still significantly correlated with functional connectivity between the left inferior frontal gyrus (LIFG) seed and the remaining dMPFC subsystem regions (*r*(75) = .24, *p* = .037) and the total conversation duration during the month after the scan day continued to not significantly correlate with the LIFG-dMPFC subsystem connectivity (*r*(71) = -.0002, *p* = .999).

### Passive Mobile Sensing

We used Wang et al.’s (2014) StudentLife iOS and Android applications to collect mobile-sensing data. Participants downloaded the app (77 iOS and 11 Android users) at the onset of the study and the application passively sensed audio and GPS features. Related publications explain the preprocessing and analyses of passively-sensed data (Chen et al., 2013; N. Lane et al., 2012; Rabbi et al., 2011; Wang et al., 2014). We maintained the temporal resolution in the order of weeks (multiples of 7 days) to remove any inconsistencies in the students’ class, work, and extracurricular schedules. In other words, any structure to conversation behavior specific to certain days of the week (e.g., sorority meetings on Thursdays; history class on Mondays) was held constant by creating week-specific values. We calculated the conversation features over two, separate 4-week (28 days ≈ 1 month) intervals, the first ending on the day before the scan and second starting on the day after the scan. For ease, throughout the manuscript we refer to these variables as “1 month” in the past and future.

### Tracking Conversation Time

Smartphone microphones were used as sensors to collect audio features with a sampling ratio of 1 min ON to 3 min OFF. The detected features were fed into a hidden Markov model (HMM) to classify the audio as human voice and to detect if the audio may resemble a conversation (N. D. Lane et al., 2014). HMM-based classifiers have been shown to have high audio-based feature-detection accuracy (84–94%; N. Lane et al., 2012; Rabbi et al., 2011). Once determined that the participant was exposed to conversations occurring in close vicinity, the microphone keeps logging the features until the detected conversation has ended. For ethical practices and in accordance with Dartmouth College’s IRB, all personal and identity-based information, including voice and content of the conversations, were not saved. All the feature-detection was carried out on-line, and only the features approved by the IRB were logged. From the logged conversation data per subject, we could extract the duration of conversation, calculated as the sum of all individual conversation durations over 1 month in the past and future (separately) per subject.

### Tracking Location

GPS coordinates were logged to track the movement of a subject on and around the campus and surrounding area. We take advantage of these GPS logs to track conversation at various locations, such as at and around eateries. The mobile application logged the GPS coordinates every 10 minutes, sampling a total of 144 location points a day per subject. This spatial data consisted of altitude (not used), latitude, longitude, and accuracy in meters, the last of which implies higher accuracy for lower values. The spatial data was collected parallel to the conversation data, and we merged these two streams of data in a rolling fashion, since a conversation log will have a location ping recorded within 5 minutes of either occurring. Observations containing more than 20 minutes of inconsistencies were dropped, resulting in an average 3% data loss (daSilva et al., 2021). Density-based spatial clustering of applications with noise (DBSCAN; Ester et al., 1996) was used to cluster the cleaned data to spatial locations on and around campus. The campus where data was collected (Dartmouth College) is located in a small town and first-year students all live on campus. Thus, constraining GPS coordinates to locations on and around campus still reflects the typical, day-to-day lifestyle of participants.

The locations were then further grouped into six categories. We chose these six categories (as opposed to additional categories) because this allowed for each category to have at least 50 participants frequent it, to help guard against unreliable estimates. More specifically, N=50 provides 80% power to detect an effect size as large as and in the same direction (i.e., one-tailed analysis) as the one reported in our primary eateries result (see below; *r* = .34). The categories are: (1) Eateries (e.g., campus cafeterias and local restaurants), (2) Classroom Buildings, (3) Cultural Venues (e.g., the campus theatre and museum), (4) Greek Housing (where individuals in the same fraternity or sorority live together and throw social events), (5) Student Housing, and (6) Libraries. For further information, refer to Supplementary Table 2.

**Figure 1:**
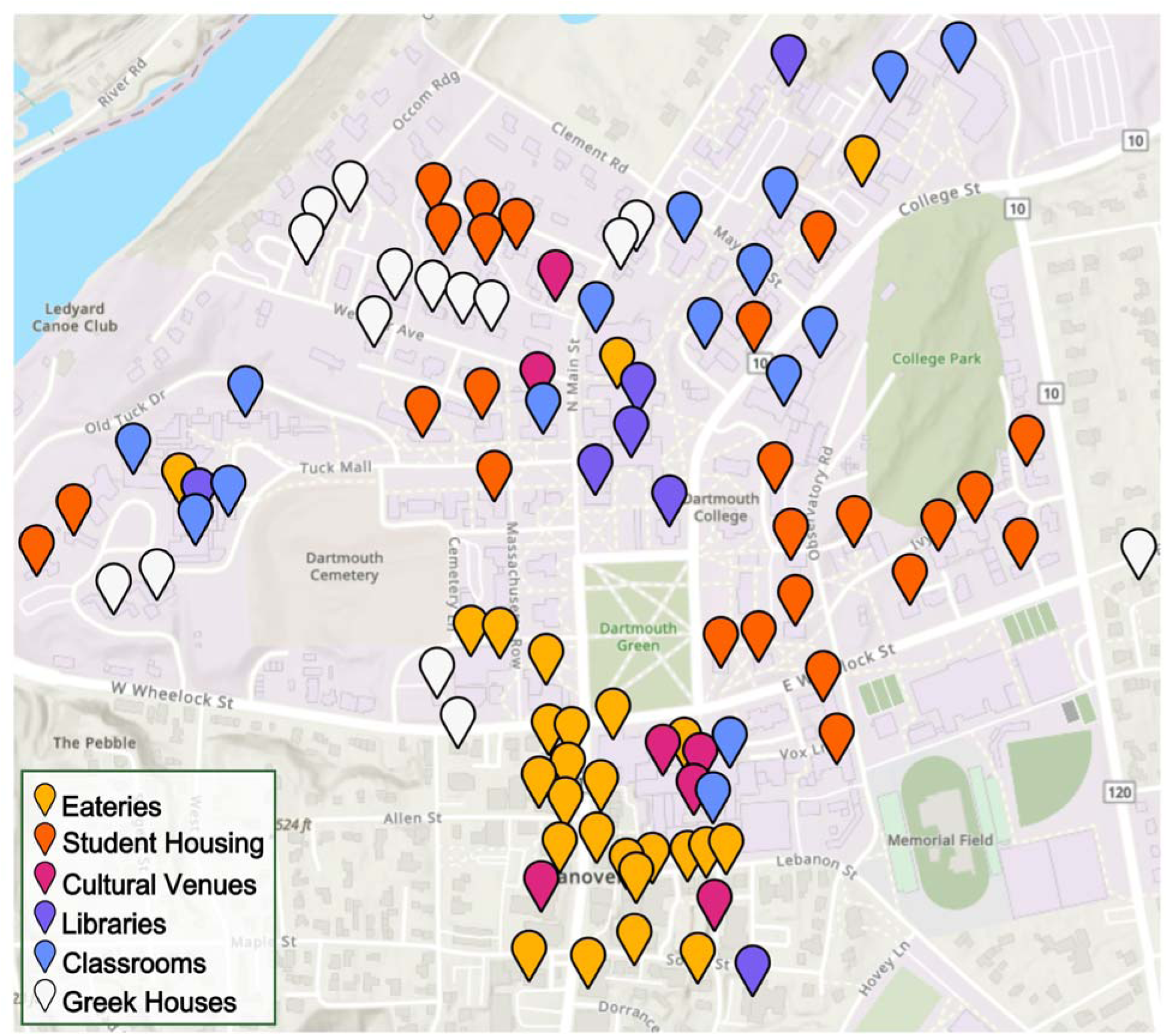
Locations used to identify in which of the six categories of locations– eateries (yellow), student housing (orange), cultural venues (magenta), libraries (violet), classrooms (blue), or greek houses (gray) –participants’ conversations took place. The map is generated using ESRI’s Python API (https://developers.arcgis.com/python/). The complete set of locations is listed in Supplementary Table 2.

### Resting State fMRI

#### fMRI Data Acquisition

Brain imaging data were collected with a 3T Siemens MAGNETOM Prisma MRI scanner with 32-channel phased array head coil. The scan session included resting state scans, as well as tasks designed to measure cognitive processes unrelated to the scope of the present research question. For this reason, we focus analyses on the resting state data.

Blood oxygen level dependent (BOLD) functional MR images were acquired using an EPI gradient echo sequence with 2.5 × 2.5 × 2.5 mm isotropic voxels, reaction time (TR) of 1000 msec and echo time (TE) of 33 ms, 3.5 mm slice thickness with 0.5 mm skip between slices, Field of View (FoV) = 240 mm × 240 mm, matrix size of 96 × 96, 90° flip angle, and a sense factor of 2. A T2-weighted structural image was acquired coplanar with the functional images (MP-RAGE; 160 sagittal slices; 1 mm × 1 mm × 1 mm voxels; 9.9 ms TR and 4.6 ms TE; 0.9-mm slice thickness; 240 mm × 240 mm FoV; 8° flip angle; sense factor of 4). The resting state scan lasted 12 minutes and 54 seconds during which time participants were instructed to lie still and let their mind wander.

#### Resting state fMRI preprocessing

Preprocessing was performed using *fMRIPrep* version *20.0.3* (Esteban et al., 2019; Esteban, Ciric, et al., 2020; Esteban, Markiewicz, DuPre, et al., 2020), which is based on *Nipype* 1.4.2 (Esteban, Markiewicz, Johnson, et al., 2020; K. Gorgolewski et al., 2011; K. J. Gorgolewski et al., 2016). Full steps applied to the data are reported in Supplementary Materials. Briefly, standard preprocessing steps were performed including realignment, normalization into MNI space, spatial smoothing (full width half maximum of 6 mm) and nuisance regressors applied to the data included six motion parameters from realignment and their temporal derivatives and quadratic terms, motion spikes, CSF and white matter, and global signal.

#### Resting state functional connectivity (RSFC)

We parceled the preprocessed BOLD signals into the default network as defined by Yeo et al. (2011)’s 17 network functional atlas. Consistent with prior work, we removed voxel-clusters with fewer than 5 voxels to minimize noise (Brietzke & Meyer, 2021; Collier & Meyer, 2020; Meyer et al., 2019). This parcellation scheme is derived from a large sample (1,000 subjects) and the default network regions are anatomically similar to those determined with task-based fMRI studies examining social cognition (Amodio & Frith, 2006; Denny et al., 2012; Saxe & Kanwisher, 2003).

The Yeo et al. (2011) atlas includes the three default network subsystems: the dMPFC subsystem, the core subsystem, and the MTL subsystem. Given that the dMPFC subsystem is the subsystem most reliably implicated in high-level social information processing, we first investigated whether time spent in conversation in eateries relates to dMPFC subsystem functional connectivity, shown in Fig.2. To determine the specificity of our results to the dMPFC subsystem, we next performed follow-up, complementary analyses with the core and MTL subsystems. Supplementary Table 1 lists the regions included in each subsystem defined by Yeo et al. (2011).

**Figure 2:**
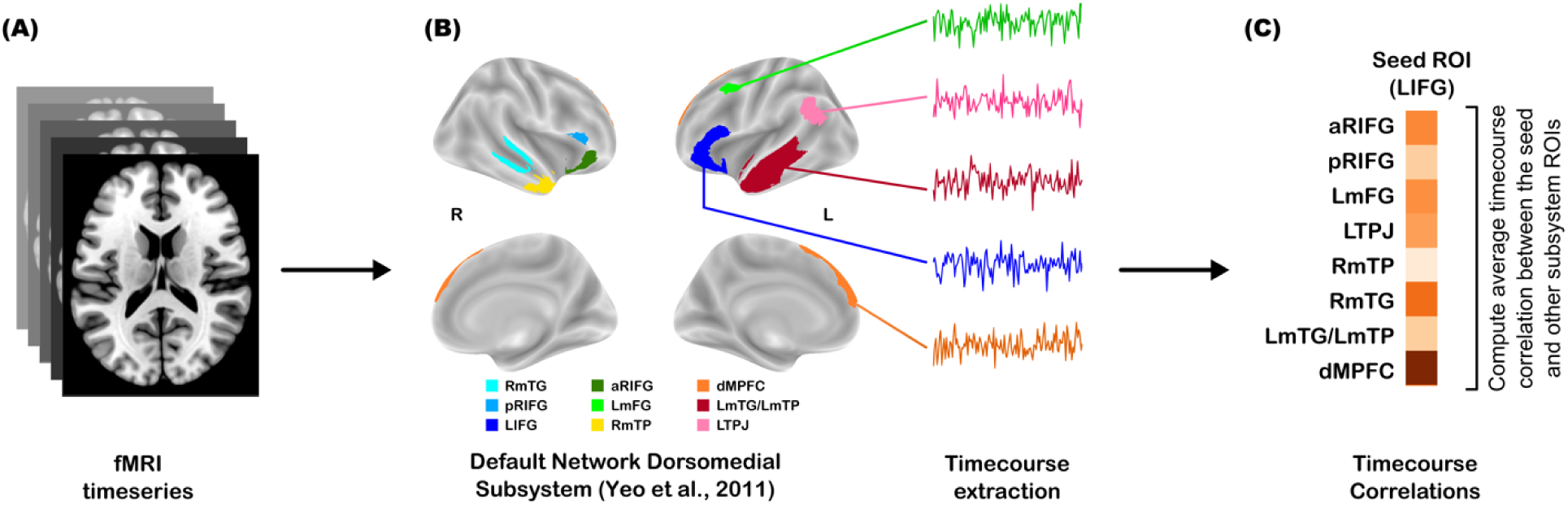
Data analysis approach for fMRI-processing. (A) Timeseries of fMRI images. (B) Parcellation of fMRI whole-brain images to the default network dorsomedial (dMPFC) subsystem, as defined by Yeo et al. (2011), followed by timecourse extraction for all the regions in the subsystem. (C) Seed-based functional connectivity (i.e., timecourse correlations) with other regions of interest (ROIs) in the dorsomedial subsystem.

Functional connectivity reflects Fisher’s z-transformed Pearson’s correlations between the timecourses of brain regions. We computed, for each subnetwork, functional connectivity in two ways. First, we performed “seeded” analyses. For each subject, we extracted the timecourse of a subnetwork region (i.e., “seed”) and determined how functionally connected (i.e., the timecourse correlations) each other region of the subnetwork is to the seed. Because this procedure corresponds with running multiple statistical tests (one for each “seed” in a subnetwork), we used Bonferroni’s family-wise error-corrected *p*-values (*p*_B_), controlling for the number of seeds in each subnetwork (dMPFC Subsystem *p*_B_(*p*=.05) = .006; Core Subsystem *p*_B_(*p*=.05) = .007; MTL Subsystem *p*_B_(*p*=.05) = .008), as a threshold to determine the statistical significance of our seeded analyses. Next, we computed the average subnetwork connectivity value for each seed. Thus, each subject had a single value reflecting how coupled subnetwork regions are to the seed. Second, we computed average, within subnetwork functional connectivity by taking the mean functional connectivity value from each non-redundant pairwise correlation. These steps were taken separately for each default network subsystem.

### Seed-based Meta-Analytic Search in NeuroSynth

We followed-up our seed-based functional connectivity analyses with voxel-wise functional connectivity analyses to further explore which portions of the most relevant seed - LIFG - drive our observed results. We conducted a meta-analytic search of peak voxel locations using NeuroSynth’s ‘Locations->Associations’ tool (Yarkoni et al., 2011). We used the following seed-based Pearson’s correlation coefficients: (1) forward-inference within-brain functional connectivity (uncorrected statistics) and (2) reverse-inference across-studies coactivation (FDR-corrected statistics) –which complement our primary seed-based analyses. The NeuroSynth voxel seeds have a default 6mm radius centered at the standard MNI-152 coordinates (with 2mm ⨉ 2mm ⨉ 2mm voxels) for each of the three peaks we observed. We then cleaned the list of associated terms to remove terms that belonged to functional regions (e.g., ‘PFC’ or ‘amygdala’), structural locations (e.g., ‘BA 45’ or ‘cortex’), terms related to other modalities (e.g., ‘heartrate’ or ‘gm volume’), incomplete or ambiguous terms (e.g., ‘hub’ or ‘implicit’), and redundant plurals (e.g. removing ‘semantics’ or ‘morals’ if ‘semantic’ or ‘moral’ is present). We made sure to retain terms that may be related to stimuli, cues, and internal processes, (e.g., positive, neutral, autobiographical, episodic, etc). This left us with 46 terms that we used for our NeuroSynth-based analyses (see Supplementary Fig.2).

## Results

### Time spent in conversation at eateries during the past month predicts dorsomedial subsystem connectivity with the left inferior frontal gyrus (LIFG) seed

Does spending time in conversations around meals relate to dMPFC subsystem function and if so, how? Given that default network functional connectivity is associated with various stable traits (Inagaki & Meyer, 2020; Vaidya & Gordon, 2013), it is possible that time spent in conversation over meals also relates to this subsystem connectivity in a stable, “trait-like” way–across both months of conversation data collection. Alternatively, based on prior work showing that real-world social experiences preferentially impact default network function (Hoekzema et al., 2022; Martínez-García et al., 2023), it is possible that past time spent in conversation around meals preferentially impacts dorsomedial subsystem connectivity. Our seed-based analyses support the latter possibility.

We found that conversation duration over the month before the scan day significantly correlated with functional connectivity between the left inferior frontal gyrus (LIFG) seed and the remaining dMPFC subsystem regions (*r*(73) = .34, *p* = .004; Fig.3A). In contrast, the total conversation duration during the month after the scan day was not significantly correlated with the LIFG-dMPFC subsystem connectivity (*r*(69) = .09, *p* = .45; Fig.3B). Next, to directly compare these correlations to one another, we carried out pairwise tests for the equality of two dependent correlations (Steiger, 1980). This demonstrated that there is a significant difference between the two brain-behavior correlation coefficients (*t*(66) = 3.2, *p* = .002). Thus, the significant LIFG-dMPFC subsystem relationship with past conversation duration is meaningfully different from the null relationship for the month following the scan. It is also notable that time spent in eateries (regardless of time spent in conversation) does not relate to LIFG-dMPFC subsystem connectivity (past month *r*(75) = .04, *p* = .71; future month *r*(69) = -.04, *p* = .74). This further suggests that time spent in conversation around meals (beyond time spent around meals more generally) relates to LIFG-dMPFC subsystem function.

To further assess the robustness of our LIFG-dMPFC subsystem results, we used bootstrapping, resampling with replacement from our sample distribution, to estimate the same relationship between functional connectivity and conversation duration with each new subsample. This approach allows us to see what correlations we would observe from independent draws from the population. Next, we performed permutation tests to determine the number of times we would observe our correlation by chance, another (more rigorous) approach to statistical testing the correlation’s significance. We ran 25,000 iterations of Pearson’s correlations over randomly permuted data to derive a corrected *p*-value. As shown in Fig.3C, the correlation between conversation duration at eateries during the prior month and LIFG-dMPFC subsystem connectivity (orange solid line) is significant and it falls outside of the null distribution of permuted *p*-values. In contrast, the correlation between LIFG-dMPFC subsystem connectivity and conversation duration at eateries the following month (blue solid line; Fig.3C) is non-significant, with the *p*-value falling within the null distribution of permuted *p*-values marked between the black, dashed lines.

**Figure 3:**
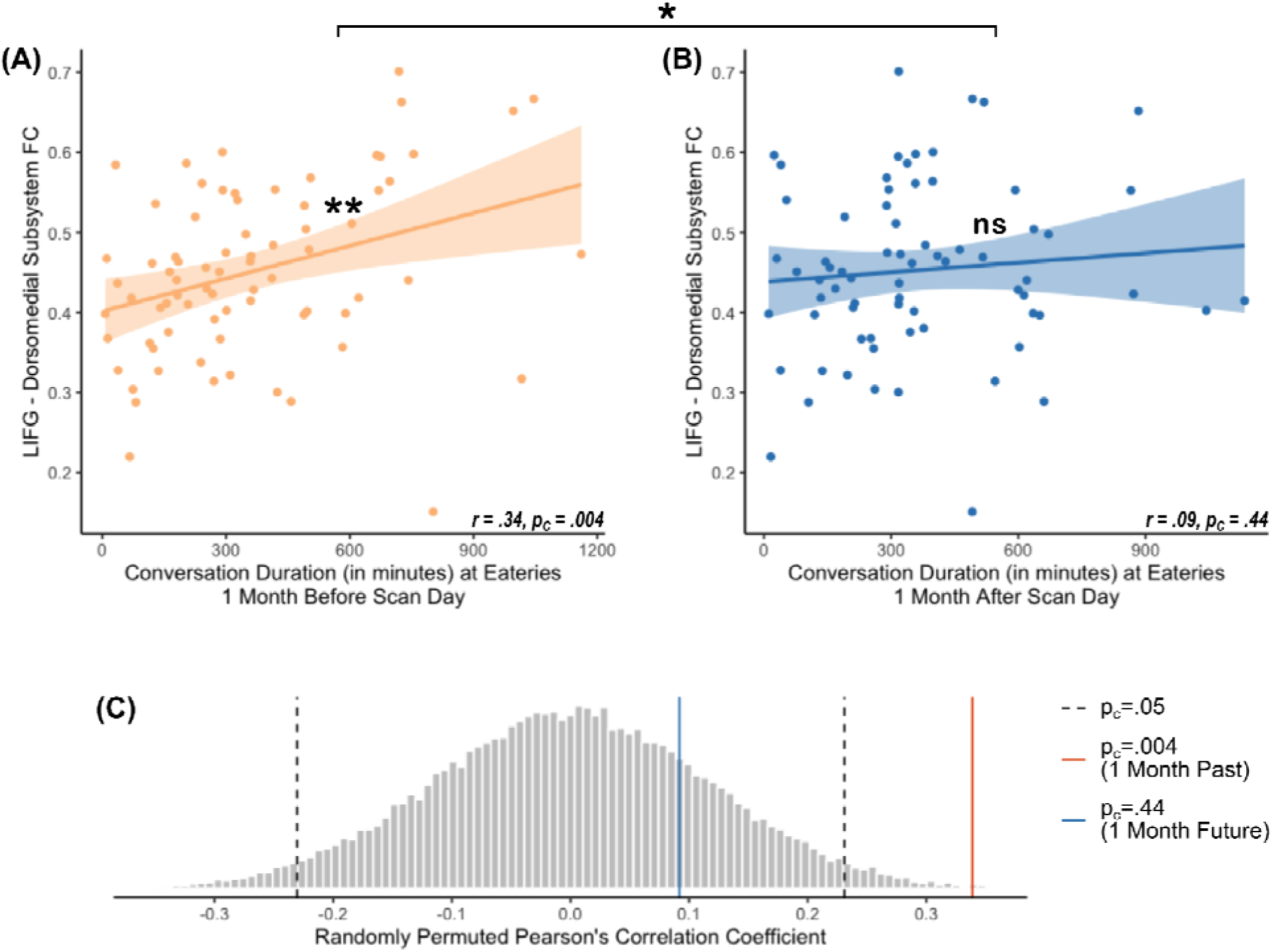
Time spent in conversation at eateries related to average LIFG-dorsomedial subsystem functional connectivity. Panels A-B present scatterplots demonstrating the relationship between time spent in conversation in the past (A) and future (B) month with LIFG-dorsomedial subsystem functional connectivity. Standard error is displayed in the bands around the best fine lines. The bracket-bar above the two scatterplots indicates a significant difference between the two correlation values. Panel (C) shows the permutation test to extract corrected p-values (symbolized as ‘p_C_’). The histogram (grey) shows the null distribution for randomly permuted Pearson’s correlation coefficients between the average functional connectivity of the Left IFG with the other dorsomedial subsystem regions and total conversation duration at eateries (in minutes) over 1 month before (orange) and 1 month after (blue) the scan day. The distributions were identical for both the permutation tests. The observed corrected Pearson’s coefficients are marked in solid-colored lines. The test for prior month’s conversations in orange (at r = .34, p_C_ = .004) is well outside the null distribution, while the future month’s test in blue is not (at r = .09, p_C_ = .44). The black, dashed line mark the coefficient value at significance (p_C_=.05).

When the other dMPFC subsystem parcels were used as seeds, the relationships between functional connectivity and conversation duration 1 month before, as well as 1 month after, the scan day were non-significant (|*r*’s| < .22, *p* > .05, 69 ≤ *df* ≤ 73). The relationship between average within-network functional connectivity for the entire dorsomedial subsystem (i.e., collapsing across all ROI pairs) and conversation duration 1 month before and after the scan day is also nonsignificant (*r*(73) = .13, *p* = .27 for the prior month; *r*(71) = -.0004, *p* = .99 for the succeeding month). Finally, we observed null results for past and future months with the Core-DMN and MTL subsystems– both with the seed-based and average subsystem functional connectivity approaches (|*r*’s| < .23, *p* > .05, 69 ≤ *df* ≤ 74) –with two exceptions in MTL-subsystem, left medial occipital lobe (*r*(72) = -.26, *p* = .03) and left fusiform gyrus (*r*(71) = -.29, *p* = .02), neither of which passes the respective Bonferroni correction (*p*_B; MTL_ < .008). Collectively, results suggest that conversations around meals may play a particularly important role in aligning dorsomedial subsystem function with the LIFG node in this subsystem.

### LIFG-dMPFC subsystem results are preferential to conversations at eateries

Is the LIFG-dMPFC subsystem result preferential to past conversations at *eateries*? If our findings reflect the impact of conversations around meals on default network brain function, then we should not see similar results for time spent in conversations at other locations. To examine this, we assessed the relationship between LIFG-dMPFC subsystem connectivity and conversation duration one month prior to the scan at other location categories. We focused here on one month prior to the scan because that is the timepoint in which conversations at eateries relate to LIFG-dMPFC connectivity; focusing on one month prior here thus helps constrain the number of follow-up tests assessing location specificity.

These analyses yielded non-significant results (|*r*’s| < .18, *p*’s > .25, 50 ≤ *df* ≤ 72; Fig.4). Additionally, significant differences in LIFG-dMPFC subsystem connectivity and duration of conversations over 1 month before the scan emerged when eateries were directly compared with the correlation to cultural venues (*t*(48) = 2.57, *p* = .017) and when compared with libraries (*t*(61) = 2.11, *p* = .045). A marginal significance was found for the comparisons between eateries and greek houses (*t*(47) = 1.75, *p* = .08), as well as t between eateries and classrooms (*t*(68) = 1.79, *p* = .08). That said, no significant differences were found when the eateries correlation was compared with student housing, athletic facilities, or marketplaces (*t*’s < 1.4, *p*’s > .10, 36 ≤ *df* ≤ 67). A detailed break-down of the location-comparison results are provided in Supplementary Table 3. It is also noteworthy that participants spent the longest proportion of time in conversations at eateries relative to every other category of locations, excluding greek houses; (see Supplementary Fig.1 for more details).

**Figure 4:**
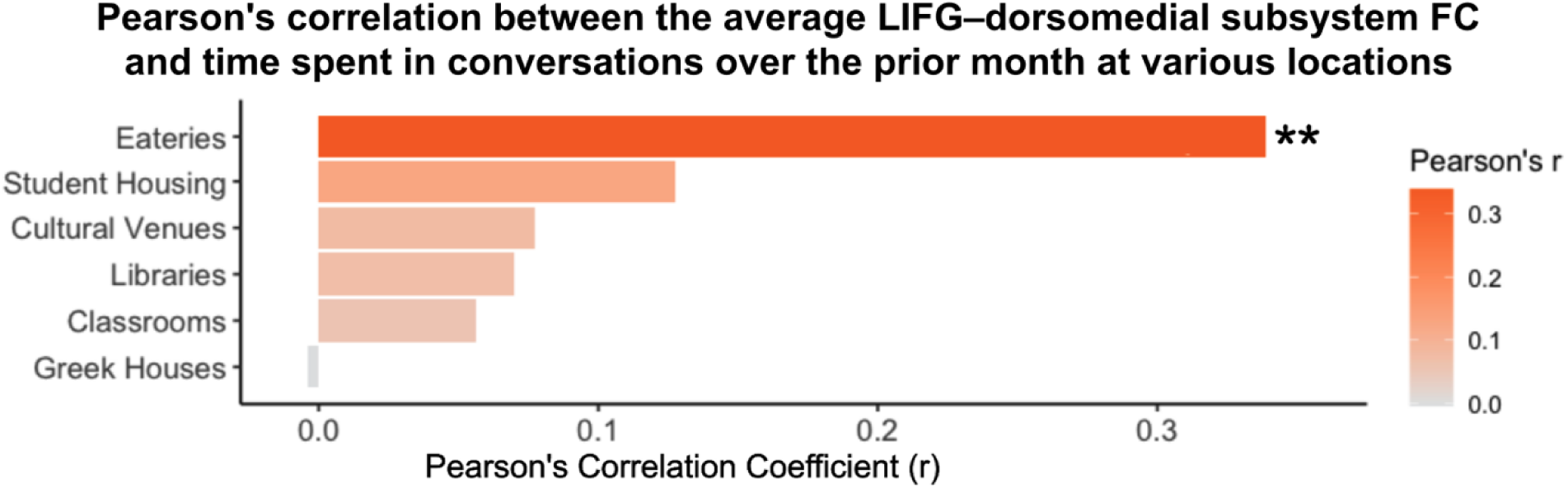
Correlations between the LIFG-dorsomedial subsystem connectivity and time spent in conversations during prior month at different locations: The bars on the plot are colored and ordered from most positive correlation coefficients (in orange) to most negative coefficients (in blue). Asterisks (**) indicate a p < .006. Note that the eateries correlation, shown in Fig.3A, is also shown here to demonstrate the specificity to eateries relative to other locations.

### Time spent in conversations at eateries impacts dMPFC subsystem connectivity with ventral LIFG

Our results indicate that more time spent around conversations at eateries impacts LIFG-dMPFC subsystem connectivity. Given that the LIFG parcel is relatively large and that the LIFG has been implicated in many cognitive and affective processes (Arioli et al., 2021; Buhle et al., 2014; Grecucci et al., 2013; Hooker et al., 2008; Klaus & Hartwigsen, 2019; Liakakis et al., 2011; Nakic et al., 2006; Satpute et al., 2014; Uddén & Bahlmann, 2012; Xu et al., 2020), we next ran exploratory analyses to see if a specific subarea of this region drives the results.

**Figure 5:**
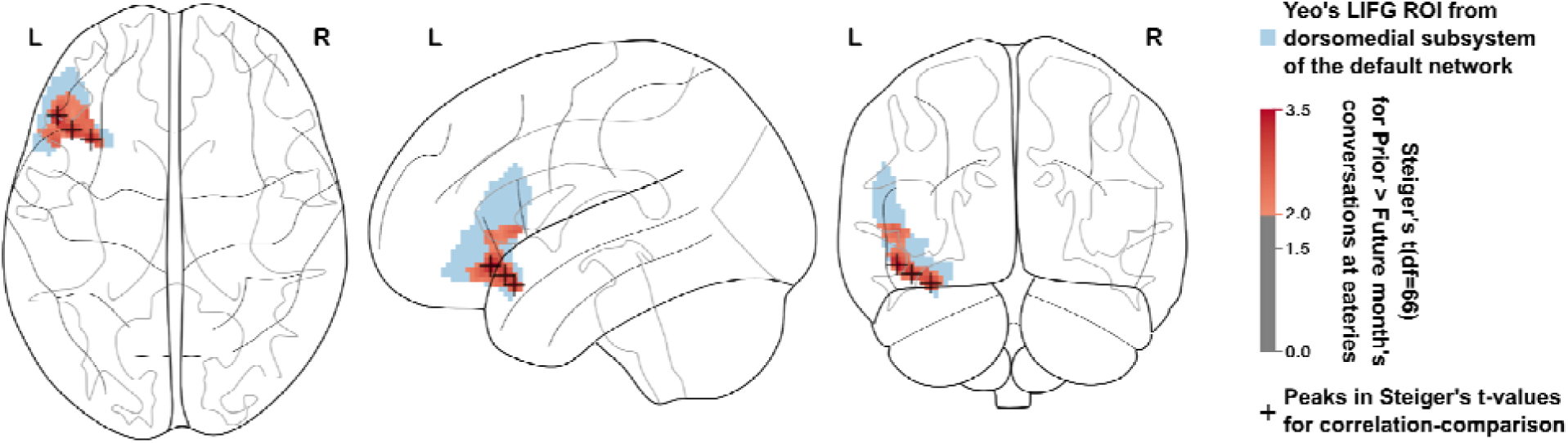
Cluster of voxels showing significant Steiger’s t-value for the functional connectivity with other dorsomedial subsystem regions when correlated to the conversations at eateries over the prior and future months: The glass-brain plots in the axial (left), sagittal (middle), and coronal (right) planes are shown. In each plane, the Left IFG mask within the dorsomedial subsystem of Default Network (according to Yeo et al., 2011) is shaded in blue) and voxel-map of significant Steiger’s t-values (t(66) > 1.99, p < .05). For each voxel’s functional connectivity within the dorsomedial subsystem, Steiger’s t-values compare the correlations with the duration of conversations occurring at eateries over the prior month to that taking place over the future month. Three ‘peaks,’ marked with ‘+,’ are found in the LIFG at: x=-48 y=28 z=12, x=-34 y=18 z=-20, and x=-42 y=22 z=-16.

We extracted voxel-wise, gaussian-smoothed time-series from the LIFG seed. We then correlated these vectors with the timeseries of other dorsomedial subsystem parcels and averaged the Fisher’s z-scored Pearson’s correlations. This approach yields LIFG voxel-dMPFC subsystem functional connectivity per subject. We then compared the correlation coefficients of these voxel-dorsomedial subsystem average connectivity variables with the respective times spent around conversations at eateries over the prior vs. future months. We then calculated Steiger’s t-values (which compare dependent correlation values) for every voxel in LIFG and threshold the *t*-values to *p* < .05; Fig.5. Our decision to use this correlation-comparison statistic follows our first set of results that not only show that past conversation duration significantly correlates with IFG-dMPFC subsystem connectivity, but further that this relationship is significantly different from the parallel relationship for the future month. The voxel-wise approach revealed three peaks, all located in the ventral portion of the LIFG (peaks at: x=-48 y=28 z=12; x=-34 y=18 z=-20; x=-42 y=22 z=-16). Examining the results for each month separately showed these voxels’ connectivity to the dorsomedial subsystem are significantly correlated with the prior month’s conversations at eateries (*r*(66)’s > .25, *p*’s < .04) and not significantly correlated to that during the future month (|*r*(66)|’s < .17, *p*’s > .10). Thus, time spent around conversations at eateries preferentially impacts functional connectivity between the dMPFC subsystem and the ventral portion of LIFG.

**Figure 6:**
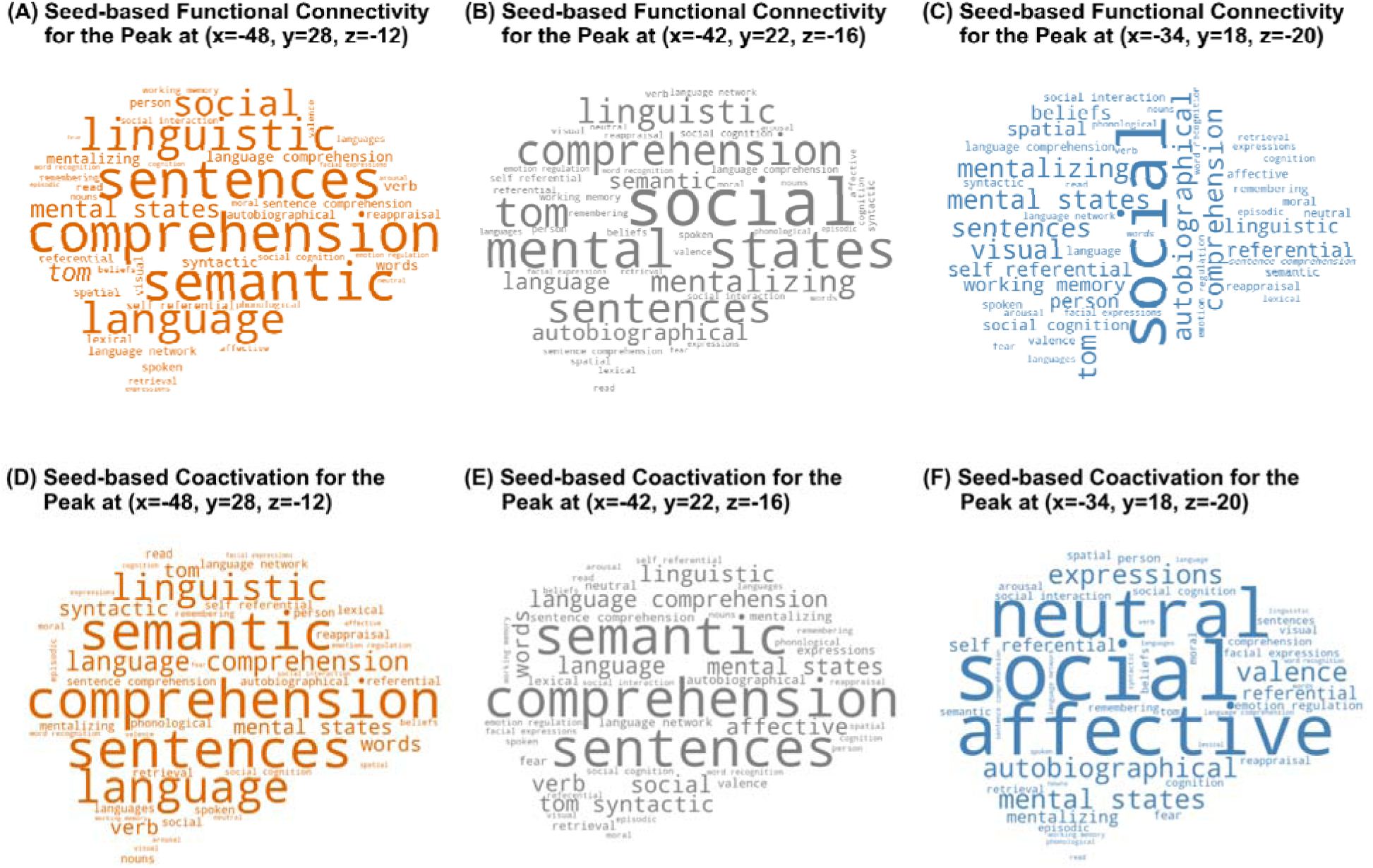
Seed-based meta-analytic search of the three peak locations: The wordclouds show the processes often implicated with the three voxel-cluster peaks in the ventral LIFG. The words are sized according to their respective Pearson’s correlation coefficients. The seeds (standard MNI-152 coordinates) are the three peaks located in ventral-LIFG (see Fig.5). In the top row (A-C), the terms used in the respective wordclouds are uncorrected Pearson’s r-values for seed-based functional connectivity during rest for locations that fall within 6 mm of the peaks found at (A) x=-48 y=28 z=-12, (B) x=-42 y=22 z=-16, and (C) x=-34 y=18 z=-20. In the bottom row (D-F), the terms used in the respective wordclouds are FDR-corrected Pearson’s r-values for seed-based meta-analytic coactivations during tasks, as per the studies in NeuroSynth’s database (Yarkoni et al., 2011), for locations that fall within 6 mm of the peaks found at (D) x=-48 y=28 z=-12, (E) x=-42 y=22 z=-16, and (F) x=-34 y=18 z=-20. For a full list of the words and their respective effect sizes for each location, see Supplementary Fig.2.

Next, we used NeuroSynth’s ‘Locations->Associations’ tool (Yarkoni et al., 2011) to conduct a seed-based meta-analytic search of the three peak locations from the ventral LIFG. The first peak, x=-48 y=28 z=-12, is strongly related to terms often associated with language comprehension– e.g., comprehension, sentences, semantic, language, etc. –for both the NeuroSynth statistical measures of functional connectivity (Fig.6A) and coactivation during tasks (Fig.6D). Similarly, we found that another peak, at x=-34 y=18 z=-20, is strongly related to terms often associated with social cognition, affect, and memory– e.g., social, mental states, affective, mentalizing, autobiographical, etc. – for both the statistical measures of functional connectivity (Fig.6C) and coactivation (Fig.6F). It is also noteworthy that the peak lying in the middle of these two peaks (at x=-42 y=22 z=-16) revealed overlapping functions with the other two peaks. For example, this middle coordinate peak is functionally connected to networks associated with other social-affective processes– showing forward-inferenced associations with terms like ‘social’ and ‘mental states’ (see Fig.6B) –and meta-analytically coactivates with terms that relate to linguistic processes (e.g., ‘comprehension’ and ‘semantic’; see Fig.6E). Moreover, we find memory-related terms coactivating for all three peaks (e.g., *r*’s ≥ 0.19 for ‘autobiographical’, *r*’s ≥ 0.14 for ‘retrieval’, *r*’s ≥ 0.1 for ‘episodic’; see Supplementary Fig.2). These analyses support the previous results. They further suggest that real-world conversations may influence how the LIFG, particularly the ventral portion, works with the rest of the dorsomedial subsystem to process social, affective, memory, and linguistic information.

## Discussion

Commensality–sharing a meal with others–is a pervasive human behavior. Anthropologists and sociologists have long suggested commensality, perhaps through the advent of fire, provided humans a dedicated time for social interaction (Wiessner, 2014). Some have even suggested that commensality played a critical role in human brain evolution (Dunbar, 2017; Powdermaker, 1932). Yet, to date, whether and how shared meals impact brain function remained unexamined. By combining methods from passive-mobile sensing and brain imaging, we identified social brain regions impacted by commensality. Undergraduate participants’ conversations and locations were unobtrusively tracked for two months. Roughly halfway through mobile sensing data collection, participants also completed a standard resting state scan, which measures endogenous brain function. Time spent in conversations around meals at cafeterias and restaurants during the prior (vs. future) month predicted greater functional connectivity in the dorsomedial default network subsystem, specifically between the LIFG and the rest of the subsystem regions. The dorsomedial subsystem is critical to high-level social inference, such as inferring mental states, retrieving social semantic knowledge, and reasoning about interpersonal relationships (Andrews-Hanna et al., 2010; Buckner et al., 1999; Collier & Meyer, 2020; Mitchell et al., 2004; Satpute et al., 2014; Saxe & Kanwisher, 2003; Van Overwalle & Baetens, 2009). Collectively, results suggest the conversations afforded by shared meals may play a key role in enhancing default network function, specifically in the dorsomedial subsystem.

The relationship between time spent around conversation at eateries and LIFG-dorsomedial subsystem connectivity was remarkably preferential to these brain regions. For example, neither the other seed regions within the dorsomedial subsystem, nor seed regions within the other default network subsystems (i.e., core; MTL) yielded meaningful results. LIFG-dorsomedial subsystem connectivity was also unrelated to time spent in conversation at other locations, such as student housing, cultural venues, and classrooms. In fact, only time spent during the past month around conversations at eateries (rather than time spent at eateries more generally) predicted LIFG-dorsomedial subsystem connectivity, suggesting the *conversations* afforded by shared meals is key.

The fact that the result was specific to time spent around conversations during the prior (vs. future) month is particularly important. This helps rule out the possibility that commensality and LIFG-dorsomedial subsystem connectivity have a “trait like” relationship. If people who tend to have a lot of conversations over meals also have greater LIFG-dorsomedial subsystem connectivity, then we would have found significant relationships for both the past and future months. Although speculative, it is possible that time spent in conversation over meals causally impacts LIFG-dorsomedial subsystem function.

This possibility compliments and extends prior work demonstrating that changes to the environment that enhance the need for social information processing change default network function. For example, in non-human primates, random assignment into larger vs. smaller social networks for an average of six months increases functional connectivity between brain regions homologous to the human default network (Sallet et al., 2011). In humans, adults who transition into parenthood (vs. not), show systematic changes to default network function (Hoekzema et al., 2022; Martínez-García et al., 2023). This effect is not wholly dependent on biological changes occurring during pregnancy, given that male adults that transition to fatherhood (vs. not) also show systematic changes to default network function (Martínez-García et al., 2023; Paternina-Die et al., 2020). The parenthood results have been interpreted as reflecting the increased social cognitive demands posed by the transition to parenthood, for example anticipating the future child’s needs and imagining the self in a new role as a parent. Conversations over meals may be another social context among many that enhance the need to think about people, in turn impacting default network function.

The results also add to our understanding of the LIFG, particularly in everyday social life. LIFG encompasses a large swath of cortex and is implicated in numerous cognitive and affective capacities. In the context of social cognition, LIFG is frequently implicated in mentalizing (Schurz et al., 2014), the process of inferring people’s mental states and traits. Lesion studies suggest the IFG may be particularly important for mentalizing about emotional states, such as determining what someone else is feeling based on their facial expressions (Shamay-Tsoory et al., 2009). Outside of social cognition research, bilateral IFG is also frequently implicated in semantic knowledge and cognitive control (Botvinick et al., 2001; Fales et al., 2008; Greene et al., 2004). Relatedly, LIFG frequently co-activates with the dorsomedial prefrontal cortex (dMPFC), a key node of the dorsomedial subsystem, during tasks involving effortful, social semantic knowledge retrieval. For example, when accessing social semantic knowledge, the LIFG effortfully retrieves social semantic knowledge from memory, while the dMPFC selects which aspect of the retrieved social knowledge is most relevant to the social task at hand (Satpute et al., 2013, 2014). Zooming out to everyday social life, prior work has also found that the size of one’s active social network on social media (Mori & Haruno, 2022) is predicted by resting state functional connectivity between the LIFG and other dorsomedial subsystem brain regions (dMPFC; TPJ). This again points to the possibility that real-world engagement in social experiences is linked to LIFG-dorsomedial subsystem function. Indeed, an exploratory, voxel-wise functional connectivity analysis of our data revealed three ventral peaks in the LIFG drive our results. We entered the peaks into NeuroSynth (Yarkoni et al., 2011), a meta-analytic platform that identifies the constructs most reliably associated with brain coordinates. The peaks were associated strongly with the terms “language”, “social”, and “affect” as well as terms related to memory such as “autobiographical” and “retrieval.” While the LIFG performs many functions, the voxel-wise results further hint at the possibilities that LIFG helps support social semantic memory retrieval, as well as emotional processes, during communication.

The IFG appears as part of the default network specifically when graph-based functional connectivity approaches are used to identify the network in resting state scans (Yeo et al., 2011). Historically, the default network gets its name from the observation that regions in the network show greater univariate (i.e., mean) activity “by default” during passive rest relative to many cognitive tasks. When the default network is defined in this way, the LIFG is not identified (Raichle et al., 2001; Shulman et al., 1997; Spunt et al., 2015), likely because the region is involved in many cognitive tasks demanding attention to external stimuli. This may make the region particularly important to navigating conversations, which require a combination of internally directed social cognition (e.g., mentalizing about our conversation partner) with externally directed cognition (e.g., making inferences about external features of a speaker, such as their facial expressions; regulating the direction of the conversation). Future work utilizing ‘hyperscanning’—in which two or more participants undergo neuroimaging simultaneously—may help unravel how communicators balance internal and external information-processing during conversation and the potential role of the LIFG in this process.

Our results add to a growing literature emphasizing the importance of naturalistic approaches in neuroscience research (Grall & Finn, 2022; Nastase et al., 2020; Wheatley et al., 2019; Zaki & Ochsner, 2009). In this literature, typically the naturalism refers to the stimuli shown during fMRI scanning. Here, we show that measuring real-world social behavior may be just as important. It is hard to imagine how the role of commensality in brain function could be assessed without a push towards naturalistically measuring real-world conversation over meals. While liquid foods such as milkshakes can be delivered to participants during neuroimaging, such an experience would hardly match the social complexity of a shared meal in everyday life. Self-reported time spent in conversation over meals also seems susceptible to retrospective errors (Collopy, 1996; Mastrandrea et al., 2015), especially since people have many misconceptions about the timing of conversations (Hirschi et al., 2023; Mastroianni et al., 2021). Meanwhile, passive mobile sensing is a growing field in its own right (Mack et al., 2021; Pillai et al., 2023; Rabbi et al., 2011; Saragosa-Harris et al., 2022; Spathis et al., 2019). Although most of this research focuses on predicting mental and physical health outcomes, mobile sensing is also a promising approach for social, cognitive, and affective neuroscientists interested in brain function. For instance, cognitive neuroscientists interested in cognitive maps could link variability in real-world spatial navigation to brain function. In our view, the results reported here, alongside a handful of other papers linking mobile sensing with brain function (Heller et al., 2020; Huckins et al., 2019; Obuchi et al., 2020), suggest this may be a fruitful approach to determining the basic neural processes tied to real-world experience.

More broadly, the results have interesting implications for understanding how social environments shape brain function. Sociologists refer to locations where individuals gather, outside of work and residence, as “third places” (Morisson, 2019; Oldenburg & Brissett, 1982). Although the definition of “third places” is broader than just eateries, research in this area finds that individuals seek out social interactions to a greater extent at cafeterias, restaurants, cafes, and pubs than other third places (Jeffres et al., 2009). A growing area of neuroscience research aims to link macro, community-level variables to individual brain function (Hatzenbuehler et al., 2022; Jorgensen et al., 2023; Weissman et al., 2023). For instance, in the United States, living in a state with anti-poverty programs attenuates the known relationship between living in poverty and reduced adolescent hippocampal volume (Weissman et al., 2023). However, to our knowledge, this area of research has not yet examined how the physical layout of a community, and the social affordances it provides, impacts brain function. Linking large neuroimaging datasets, like the Human Connectome Project, Adolescent Brain and Cognitive Development (ABCD) Study, and UK Biobank Study, with physical social structures like third places in a community may yield interesting links between brain and society.

### Limitations

Two limitations constrain the specificity and generalizability of our results. First, in terms of specificity, it is unclear if it is the amount of time a participant is talking, listening, or some combination that drives the results. The StudentLife app used here measures time spent around face-to-face conversation but does not identify who is speaking. The app also does not measure the content shared in conversation. Additional research is needed to determine the precise aspects of conversation necessary to influence IFG-dorsomedial subsystem connectivity. Second, the sample consists of freshman in a remote college setting. The restricted age range and minimal diversity of our sample make it hard to know if the results generalize to other communities. College freshmen are also tasked with forming an entirely new social network, which may amplify their interest in forging social connection. Whether this motivation is necessary for shared meals to influence dorsomedial subsystem function remains to be determined.

## Conclusion

Anthropologists and sociologists have argued that commensality is a human universal that may have driven human brain evolution. Yet, to date, how shared meals impact brain function remained unclear. We found initial evidence that shared meals may impact social brain mechanisms through the conversations they afford. More time spent in conversation during the prior (vs. future) month predicted greater resting state functional connectivity between the LIFG and the remaining dorsomedial subsystem of the brain’s default network. Conversations over meals may not only provide sustenance; they may also help us navigate social life.

## Acknowledgements

This work was supported by an NIMH R01 awarded to Meghan Meyer as well as an NIMH R01 awarded to Andrew Campbell.

