## Supplementary Materials for "Time spent in conversation over meals predicts default network function: Evidence from a passive mobile-sensing and fMRI study"

### Supplementary Methods

#### ***Resting state data preprocessing***

Results included in the manuscript come from preprocessing performed using *fMRIPrep* version *20.0.3* (Esteban et al., 2019; Esteban, Ciric, et al., 2020; Esteban, Markiewicz, Johnson, et al., 2020), which is based on *Nipype* 1.4.2 (Esteban, Markiewicz, DuPre, et al., 2020; Esteban, Markiewicz, Johnson, et al., 2020; K. Gorgolewski et al., 2011; K. J. Gorgolewski et al., 2016). As the creators of fMRIprep recommended, the functional preprocessing steps are reported below verbatim from the software output.

The following preprocessing was performed for the subjects’ resting state BOLD runs. First, a reference volume and its skull-stripped version were generated using a custom methodology of fMRIPrep. A deformation field to correct for susceptibility distortions was estimated based on fMRIPrep’s fieldmap-less approach. The deformation field is that resulting from co-registering the BOLD reference to the same-subject T1w-reference with its intensity inverted (Huntenburg, 2014; Wang et al., 2017). Registration is performed with *antsRegistration* (ANTs 2.2.0), and the process regularized by constraining deformation to be nonzero only along the phase-encoding direction, and modulated with an average fieldmap template (Treiber et al., 2016). Based on the estimated susceptibility distortion, a corrected EPI (echo-planar imaging) reference was calculated for a more accurate co-registration with the anatomical reference. The BOLD reference was then co-registered to the T1w reference using *bbregister* (FreeSurfer) which implements boundary-based registration (Greve & Fischl, 2009). Co-registration was configured with six degrees of freedom. Head-motion parameters with respect to the BOLD reference (transformation matrices, and six corresponding rotation and translation parameters) are estimated before any spatiotemporal filtering using *mcflirt* (FSL 5.0.9; Jenkinson et al., 2002). The BOLD time-series were resampled onto the following surfaces (FreeSurfer reconstruction nomenclature): fsaverage. The BOLD time-series (including slice-timing correction when applied) were resampled onto their original, native space by applying a single, composite transform to correct for head-motion and susceptibility distortions. These resampled BOLD time-series will be referred to as preprocessed BOLD in original space, or just preprocessed BOLD. The BOLD time-series were resampled into standard space, generating a preprocessed BOLD run in MNI152NLin2009cAsym space. First, a reference volume and its skull-stripped version were generated using a custom methodology of fMRIPrep. Grayordinates files (Glasser et al., 2013) containing 91k samples were also generated using the highest-resolution *fsaverage* as intermediate standardized surface space. Several confounding time-series were calculated based on the preprocessed BOLD: framewise displacement (FD), DVARS and three region-wise global signals. FD and DVARS are calculated for each functional run, both using their implementations in Nipype (following the definitions by Power et al., 2014). The three global signals are extracted within the CSF, the WM, and the whole-brain masks. Additionally, a set of physiological regressors were extracted to allow for component-based noise correction (CompCor; Behzadi et al., 2007). Principal components are estimated after high-pass filtering the preprocessed BOLD time-series (using a discrete cosine filter with 128s cut-off) for the two CompCor variants: temporal (tCompCor) and anatomical (aCompCor). tCompCor components are then calculated from the top 5% variable voxels within a mask covering the subcortical regions. This subcortical mask is obtained by heavily eroding the brain mask, which ensures it does not include cortical GM regions. For aCompCor, components are calculated within the intersection of the aforementioned mask and the union of CSF and WM masks calculated in T1w space, after their projection to the native space of each functional run (using the inverse BOLD-to-T1w transformation). Components are also calculated separately within the WM and CSF masks. For each CompCor decomposition, the k components with the largest singular values are retained, such that the retained components’ time series are sufficient to explain 50 percent of variance across the nuisance mask (CSF, WM, combined, or temporal). The remaining components are dropped from consideration. The head-motion estimates calculated in the correction step were also placed within the corresponding confounds file. The confound time series derived from head motion estimates and global signals were expanded with the inclusion of temporal derivatives and quadratic terms for each (Satterthwaite et al., 2013). Frames that exceeded a threshold of 0.25 mm FD or 1.05 standardised DVARS were annotated as motion outliers. All resamplings can be performed with a single interpolation step by composing all the pertinent transformations (i.e. head-motion transform matrices, susceptibility distortion correction when available, and co-registrations to anatomical and output spaces). Gridded (volumetric) resamplings were performed using *antsApplyTransforms* (ANTs), configured with Lanczos interpolation to minimize the smoothing effects of other kernels (Lanczos, 1964). Non-gridded (surface) resamplings were performed using *mri_vol2surf* (FreeSurfer).

Before analyzing the data, spatial smoothing was applied using a 6mm full-width, half-maximum Gaussian kernel. A GLM using *nltools* (Chang et al., 2020), a fMRI-preprocessing library for python 3, also regressed out additional variances associated with the global signals and motion spikes. The residuals from this linear regression were used for subsequent analyses.

### Supplementary Tables

**Supplementary Table 1: Regions in the three DMN subsystems:** *The centroids (Yeo et al., 2011) are listed in the standard MNI-152 functional atlas with 2mm×2mm×2mm voxels. ROIs with 5 voxels or less have been removed.*

| **Network** | **Centroid (MNI-152)** | | |  | **Larger Anatomical Region** | **Voxels** |
| --- | --- | --- | --- | --- | --- | --- |
|  | **X** | **Y** | **Z** |  |  |  |
| dMPFC Subsystem | 55.6 | 24.6 | 8.2 |  | R Inferior Frontal Gyrus (posterior) | 98 |
|  | 61.8 | -25.8 | -6.2 |  | R middle Temporal Gyrus | 183 |
|  | -40.4 | 12.6 | 50.4 |  | L middle Frontal Gyrus | 241 |
|  | 43 | 29.6 | -13.8 |  | R Inferior Frontal Gyrus (anterior) | 282 |
|  | -52.4 | -55.6 | 28.6 |  | L Temporo-Parietal Junction | 423 |
|  | 51.8 | 3.8 | -30.2 |  | R Medial Temporal Pole | 424 |
|  | -46.6 | 26.8 | -2.2 |  | L Inferior Frontal Gyrus | 1201 |
|  | -56.6 | -12.2 | -19 |  | L middle Temporal Gyrus / Pole | 1667 |
|  | -2.4 | 45.6 | 41.2 |  | dorsoMedial Prefrontal Cortex | 1783 |
| Core Subsystem | 61.8 | -6.8 | -18.2 |  | R Middle Temporal Gyrus | 232 |
|  | 23.4 | -48.8 | 44 |  | R Superior Frontal Gyrus | 352 |
|  | -22.4 | 28.8 | 46.6 |  | L middle Frontal Gyrus / L Superior Frontal Gyrus | 377 |
|  | -44.4 | -68.2 | 37.4 |  | L Angular Gyrus | 570 |
|  | 51.4 | -57.4 | 29.6 |  | R Angular Gyrus / R middle Temporal Gyrus | 622 |
|  | -0.6 | -50.8 | 31 |  | R / L PCC | 1494 |
|  | 0.6 | 50 | 5.2 |  | MPFC / ACC | 2648 |
| MTL Subsystem | -39 | -80 | 32 |  | L middle Occipital Lobule | 82 |
|  | 48.2 | -70.8 | 27.2 |  | R middle Occipital Gyrus | 111 |
|  | -12.4 | -55.4 | 12.6 |  | L Calcarine Gyrus / L Precuneus | 119 |
|  | 13.8 | -53.2 | 13.8 |  | R Cuneus / R Precuneus | 141 |
|  | 27.4 | -28 | -20.2 |  | R Fusiform Gyrus | 189 |
|  | -26.8 | -33 | -18 |  | L Fusiform Gyrus | 271 |

**Supplementary Table 2: Spatial clustering of tracked locations:** *Each cluster lists the locations on or around the campus. The column, ‘*Locations,*’ also includes places found outside the town of Hanover (NH-US) that students often visit. Locations with an asterisk (*) next to the cluster name have sample sizes that account for .80 power (N>50).*

| **Location Cluster** | **Locations** |
| --- | --- |
| Athletic Facilities | 'berry_sports_center', 'canoe', 'chase-field', 'football', 'ladyard_cacoe', 'leverone', 'lodge', 'softballfield', 'sport-venues', 'sport-venues-press', 'tennis', 'thompson_arena' |
| Classroom Buildings***** | 'Hillel', 'Mckenzie', 'Tuck_hall', 'batrlett', 'buchanan', 'burke', 'butterfield', 'byrnehall', 'carpenterhall', 'chasehall', 'cummings', 'currier', 'hallgarten', 'hopkins', 'hopkins-spaulding', 'kemeny', 'lsb', 'maclean', 'massrow', 'mclaughlin', 'moore', 'morano', 'murdough', 'raven-house', 'remsen', 'ripley', 'robinson', 'rockefeller-center', 'rockefeller-social-sciences', 'russell-sage', 'silsby-rocky', 'smith', 'steele', 'streeter', 'sudikoff', 'thayer_secure', 'thornton', 'vail', 'webster_cottage', 'websterhall', 'wentworth', 'whittemore', 'wilson', 'woodburyhall', 'woodward' |
| Cultural Venues***** | 'hood', 'hopkins', 'hopkins-spaulding', 'lodge', 'native_american_house', 'rockefeller-center', 'rockefeller-social-sciences', 'sphinx', 'vac' |
| Eateries***** | '53_commons', 'candela', 'capizza', 'collis', 'domino', 'hopkins-food', 'HanoverInn', 'indian_food', 'mexican-food', 'mollys', 'murphy', 'noodle', 'orient', 'pine', 'ramunto', 'salt-hill', 'starbucks', 'sushiya', 'tuktuk', 'umpleby' |
| Greek Housing***** | 'ACA', 'AD', 'AP', 'AT', 'AXD', 'BAO', 'BG', 'CGE', 'CH', 'DDD', 'EKT', 'GDC', 'KD', 'KDE', 'KKG', 'KKK', 'PDA', 'PT', 'SAE', 'SN', 'SPE', 'TDC', 'ZP', 'tabard' |
| The Green | ‘green’ |
| Student Housing***** | '13-EWL', '19-EWL', '9-EWL', 'Cohen', 'andres', 'bissell', 'brown_hall', 'channing-cox', 'east-wheelock', 'fahey-mclane', 'fairchild', 'fayerweather', 'fayerweather-south', 'french', 'gile', 'judge', 'lacasa', 'ledyard', 'little_hall', 'lord', 'maxwell', 'mcCulloch', 'morton', 'native_american_house', 'north-park', 'richardson', 'six-south', 'south-house', 'tllc', 'topliff', 'wheeler', 'zimmerman' |
| Libraries***** | 'baker-berry', 'dana-library', 'feldberg_library', 'library-default-services', 'lsl', 'sanborn' |
| Marketplace | 'NuggetTheaters', 'carson-tech_services', 'post-office', 'bookstore', 'fairbanks', 'coop', 'lemon_gift', 'Jcrew', 'talbots' |
| Medical Facilities | 'CVS', 'DHMC', 'hitchcock', 'ropeferry' |
| Religious Places | 'StDenisCatholicChurch', 'StThomasEpiscopalChurch', 'aquinas', 'christian_reading', 'church', 'rollins-chapel' |
| Miscellaneous***** | 'blunt_alumni_center', 'dartmouth_hall', 'den', 'dewey', 'mcnutt', 'ovis', 'parkhurst', 'payroll', 'reed', 'remote_offices_HREAP' |

**Supplementary Table 3: Comparing correlations of the LIFG average functional connectivity within the dorsomedial subsystem and the total conversation duration over 1 month before the scan day between that at eateries and other locations:** *Pairwise correlation comparison (Steiger, 1980) of the average LIFG functional connectivity and the total conversation duration over the month before the scan day between eateries and other locations. The corrected p-values for Pearson’s correlation (r) are calculated over a 25,000-iteration permutation test. The p-values are 2-tailed and based on t-values for a given degree of freedom (df). The significance levels assigned here are ‘.’ for marginal significance (.1 > p ≥ .05) and ‘*’ for stronger significance (.05 > p ≥ .01).*

| **Location** | **Correlation with LIFG-avgFC** | | | **Comparing correlation with eateries** | | | |
| --- | --- | --- | --- | --- | --- | --- | --- |
|  | **r** | **n** | **p** | **t** | **df** | **p (t, df)** | |
| Eateries | .34 | 75 | .004 | - | - | - |  |
| Classroom Buildings | .05 | 74 | .633 | 1.79 | 72 | .081 | **.** |
| Cultural Venues | .08 | 52 | .578 | 2.57 | 51 | .017 | ***** |
| Greek Housing | -.004 | 53 | .982 | 1.75 | 50 | .087 | **.** |
| Student Housing | .13 | 72 | .291 | 1.38 | 69 | .153 |  |
| Libraries | .07 | 68 | .577 | 2.11 | 64 | .045 | ***** |

### Supplementary Figures

**Supplementary Figure 1: Behavioral differences across different locations:** *The plots show the distribution of three behaviors – (A) conversation density, calculated as a ratio of the total duration of conversation at a location to the time spent at said location, (B) total time spent at a given location over the past month, and (C) total duration of conversation at a given location over the same period. * indicates p<.05; ** indicates p<.001; **** indicates p<.0001*

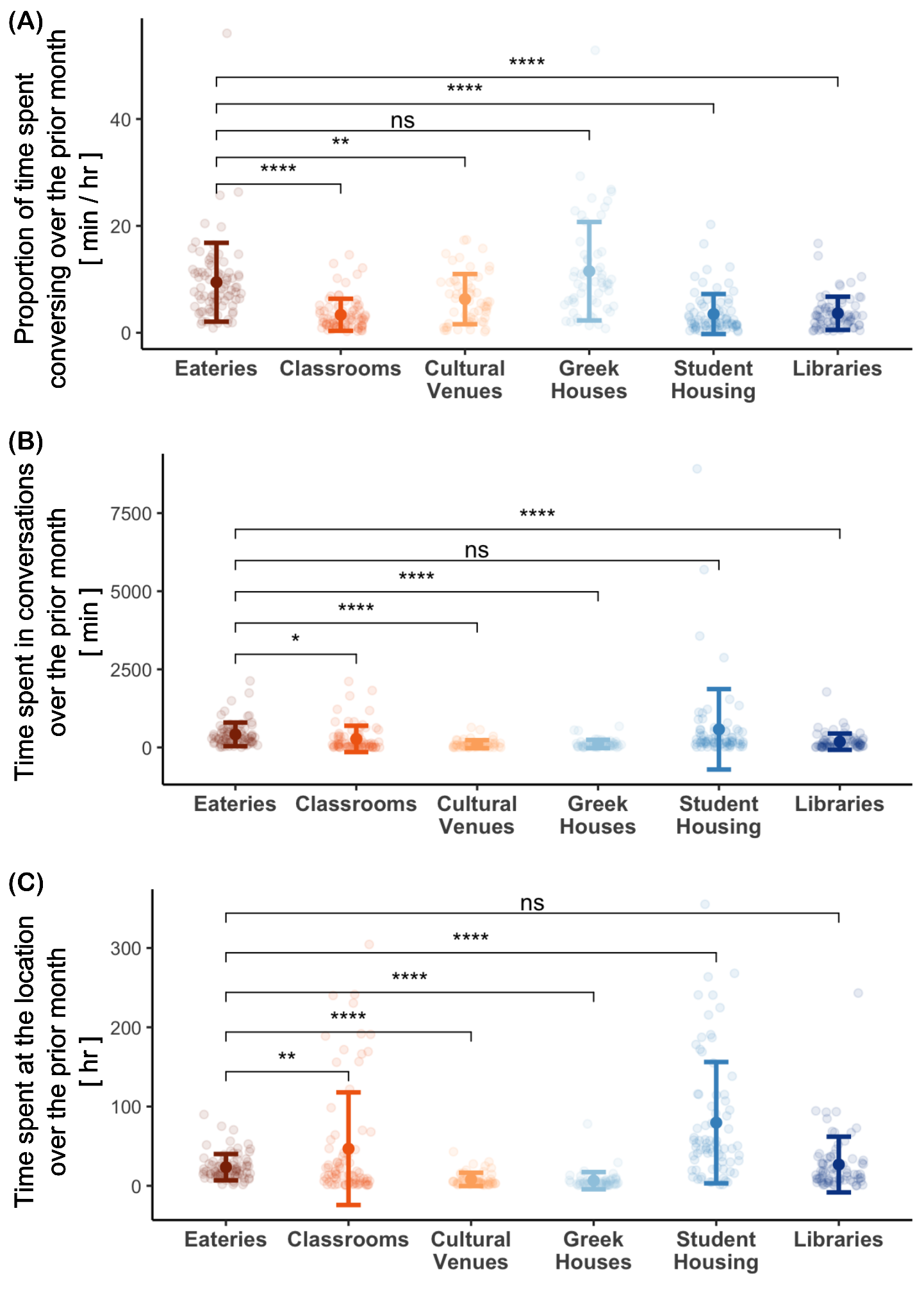

**Supplementary Figure 2: Seed-based meta-analytic search of the three peak locations:** *The bar plots show Pearson’s r-values for seed-based uncorrected functional connectivity (during resting-state; A, C, E) or FDR-corrected meta-analytic coactivations across the studies (in NeuroSynth’s database; B, D, F) with terms often implicated with the activated networks or regions (Buckner et al., 2011; Choi et al., 2012; Yarkoni et al., 2011; Yeo et al., 2011). The seed voxels (standard MNI-152 coordinates) are the three voxel-cluster peaks in ventral-LIFG. (A) Pearson’s correlation coefficient (uncorrected; on the X-axis) for the terms often implicated with resting-state networks (on the Y-axis) of functional connectivity to locations within 6mm of the seed at x=-34 y=18 z=-20. (B) Pearson’s correlation coefficient (FDR-corrected; on the X-axis) for the terms implicated from the reverse-inference map of meta-analytic coactivations across the studies (on the Y-axis) that fall within 6mm of the seed at x=-34 y=18 z=-20. (C) Pearson’s correlation coefficient (uncorrected; on the X-axis) for the terms often implicated with resting-state networks (on the Y-axis) of functional connectivity to locations within 6mm of the seed at x=-42 y=22 z=-16. (D) Pearson’s correlation coefficient (FDR-corrected; on the X-axis) for the terms implicated from the reverse-inference map of meta-analytic coactivations across the studies (on the Y-axis) that fall within 6mm of the seed at x=-42 y=22 z=-16. (E) Pearson’s correlation coefficient (uncorrected; on the X-axis) for the terms often implicated with resting-state networks (on the Y-axis) of functional connectivity to locations within 6mm of the seed at x=-48 y=28 z=12. (F) Pearson’s correlation coefficient (FDR-corrected; on the X-axis) for the terms implicated from the reverse-inference map of meta-analytic coactivations across the studies (on the Y-axis) that fall within 6mm of the seed at x=-48 y=28 z=12.*

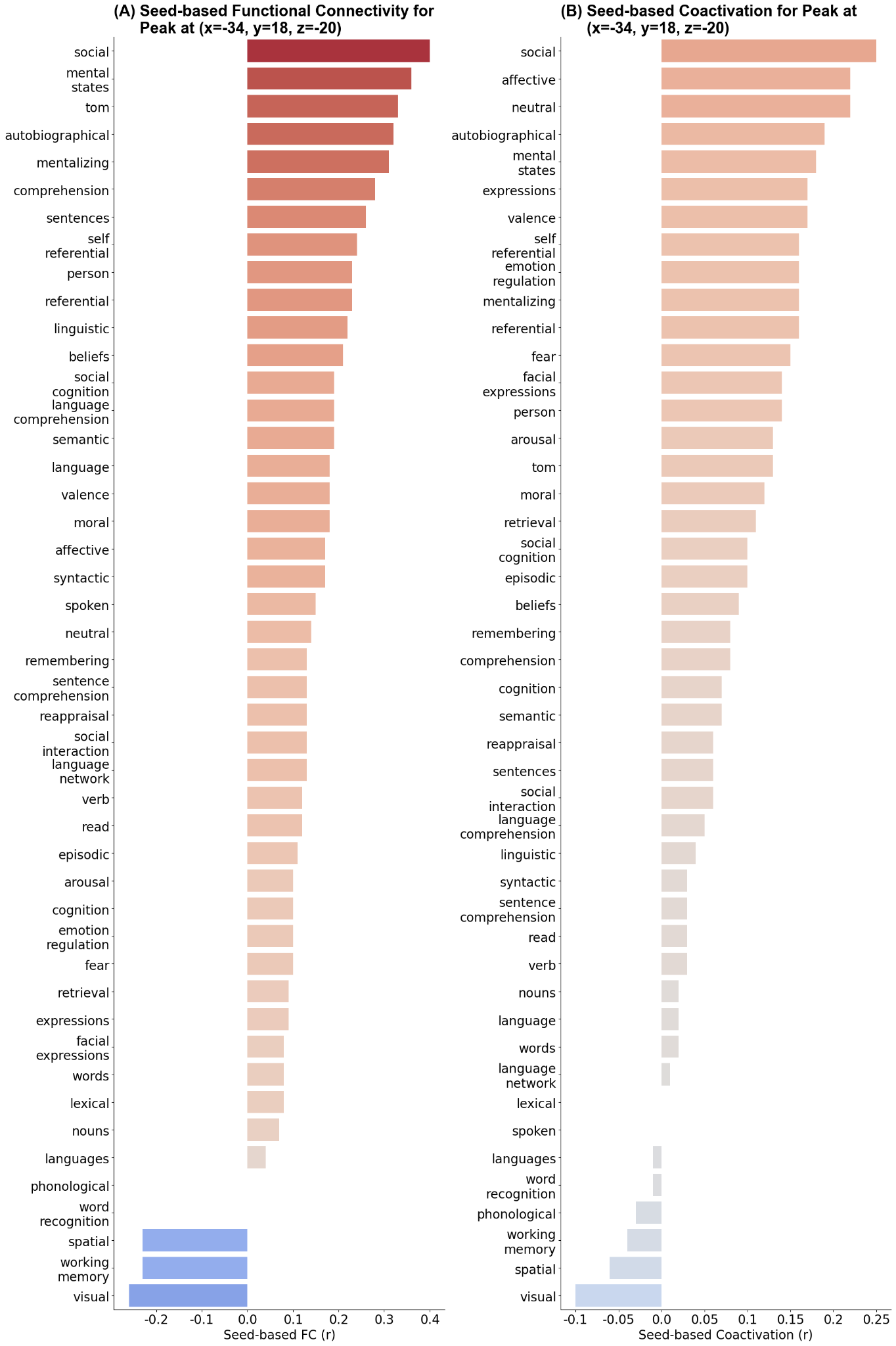

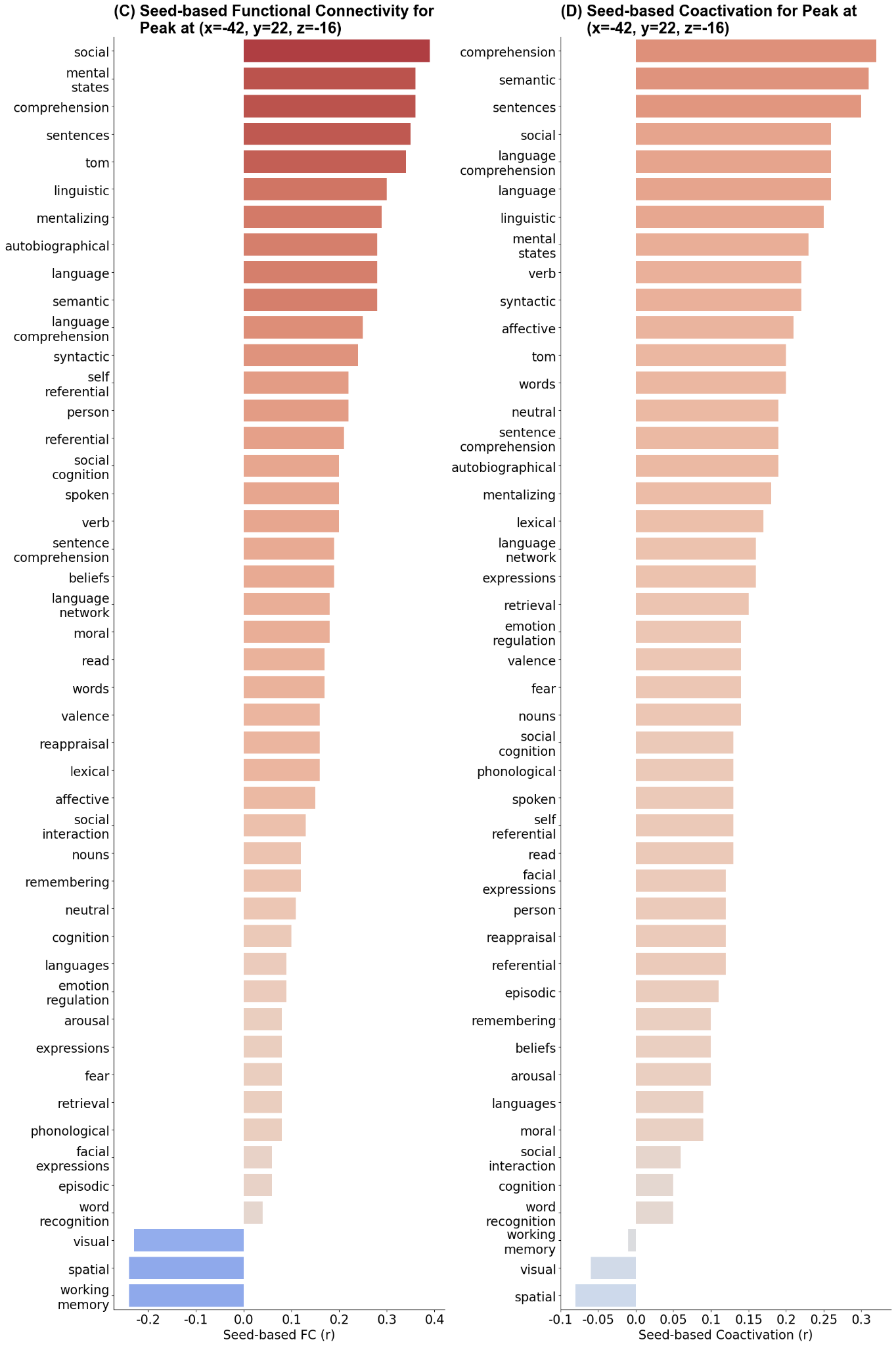

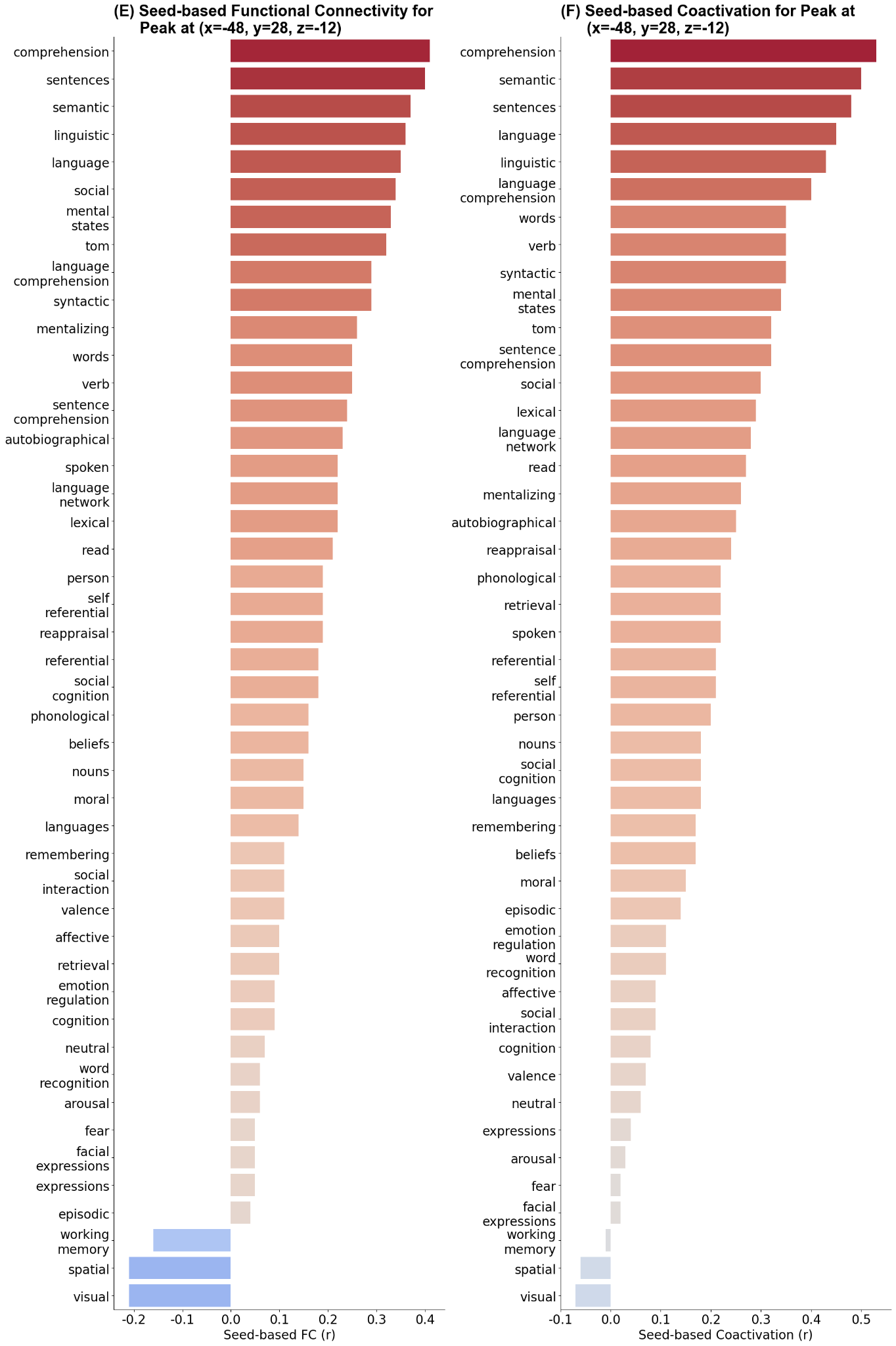
